# A Preliminary Occurrence of Extended-Spectrum and AmpC Beta-Lactamases in Clinical Isolates of Enteropathogenic *Escherichia coli* in Najaf, Iraq

**DOI:** 10.1101/512731

**Authors:** Hashim Ali Abdulameer Alsherees, Sumaya Najim Abed Ali

## Abstract

Extended-spectrum β-lactamase (ESBL) and AmpC β-lactamase producing enteropathogenic *Escherichia coli* (EPEC) have become an important clinical problem due to their resistance to multiple antibiotics. The purpose of this study was to evaluate the multi-drug resistant (MDR) and β-lactamases producing EPEC isolated from children with diarrhea. Twenty two EPEC strains were gathered from another study performed from September to November 2009. Antibiotic susceptibilities against 20 antibiotics determined by the agar disk diffusion according to the protocol recommended by Clinical and Laboratory Standards Institute guidelines (CLSI). Phenotypic confirmatory tests carried out for screening of ESBLs and AmpC β-lactamase. The isolates subjected to PCR assays with specific primers for *bla*-genes (*bla*_SHV_, *bla*_TEM_, *bla*_CTX-M_ and *bla*_OXA_, *bla*_PER_, *bla*_VEB_ and *bla*_GES_). Only 22 (3.4%) EPEC isolates recovered from clinical infections. MDR rate in EPEC was 90.9%. Phenotypic confirmatory tests showed that 13.6% and 20.0% of isolates were ESBL and AmpC β-lactamase producers, respectively. Among the EPEC isolates, percent recovery of *bla*_SHV_, *bla*_TEM_, *bla*_CTX-M_ and *bla*_OXA_ genes was 100%, 81.8%, 77.3% and 9.1%, respectively. Only 2 (9.1%) isolates had *bla*_AmpC_ gene and none of the isolates carried *bla*_PER_, *bla*_VEB_ and *bla*_GES_ genes. The results of our study showed that the ESBLs were common among EPEC isolates. Such high dissemination of ESBLs is a serious problem for public health and therefore, it is necessary to seek a program for monitoring ESBLs in Najaf hospitals. This study is the first report on the presence of the *bla*-genes in EPEC isolates in Najaf.

## Introduction

Enteropathogenic *E.coli* is an important cause of infantile diarrhea worldwide and particularly in developing countries ^[1]^. There has been an alarming increase in drug-resistant strains of EPEC in developing as well as developed countries. Several cases of antimicrobial resistance in EPEC been observed in different parts of the world ^[2]^.

Extended-spectrum beta-lactamases [ESBLs) are the enzymes that hydrolyze a wide variety of β-lactam antibiotics including oxyimino-cephalosporins and monobactams but not cephamycins, and are usually inhibited by clavulanic acid. These ESBLs are derivatives, predominantly, of class A [e.g., TEM, SHV, CTX-M, and VEB families). These enzymes, encoded by genes that are typically plasmid borne ^[3]^.

AmpC-mediated β-lactam resistance in *E. coli* is an emerging problem. High level AmpC production is typically associated with in vitro resistance to all β-lactam antibiotics except for carbapenems and cefepime. Genes for these β-lactamases found on the chromosomes of some members of the family *Enterobacteriaceae* ^[4]^. Plasmid-mediated AmpC β-lactamase has arisen through the transfer of chromosomal genes for the inducible AmpC β-lactamases onto plasmids ^[5]^. Plasmids with these genes can spread among other members of the family Enterobacteriaceae, been documented in many countries and can cause nosocomial outbreaks ^[6]^. Because of inappropriate usage of antibiotic in treatment of infection caused by ESBL producing pathogens, it seems that studies about correct detection and antibiotic resistance patterns of these organisms are necessary. However, there is no information regarding the molecular studies of ESBLs or the occurrence of AmpC β-lactamases-producing EPEC isolates in Najaf, Iraq. For this reason, this study designed to estimate the presence of ESBLs and AmpC β-lactamases in EPEC isolates.

## Materials and Methods

### Bacterial strains

Twenty two EPEC strains were gathered from another study performed from September to November 2009, that a total of 656 stool specimens from children younger than two years old with diarrhea were analyzed. *E. coli* isolates serotypically identified with EPEC polyvalent and monovalent antisera as previously described ^[7]^.

### Antibiotic susceptibility tests

All strains were characterized by antimicrobial susceptibility testing for 20 antibiotics by disk diffusion method in accordance with the Clinical and Laboratory Standards Institute guidelines^[8]^. The following antibiotics (Himedia, India) were used: amoxicillin (25 μg), amoxycillin/clavulanic acid (20 μg/10 μg), amikacin (30 μg), aztreonam (30 μg), carbenicillin (100 μg), ciprofloxacin (5 μg), cefotaxime (30 μg), ceftazidime (30 μg), ceftriaxone (30 μg), cefepime (30 μg), cefoxitin (30 μg), gentamicin (10 μg), imipenem (10 μg), levofloxacin (5 μg), meropenem (10 μg), piperacillin (100 μg), tobramycin (10 μg), ticarcillin (75 μg), trimethoprim (5 μg) and tetracyclin (30 μg).

ESBL production detected by double-disk synergy test (DDST). Disks containing ceftazidime, cefotaxime, ceftriaxone and aztreonam respectively, placed 20 mm (center to center of the disks) from the amoxicillin-clavulanic acid disk. Inoculated plates incubated aerobically at 37°C for 18 h. After incubation, distortion of the inhibition zone of any of the antibiotics towards the disc containing clavulanic acid regarded as a phenotypic screening of the presence of ESBL ^[9]^. ESBL production confirmed by both cefotaxime and ceftazidime alone and in combination with clavulanic acid. AmpC enzyme production tested by a modified three-dimension test (MTDT) as described by Coudron *et al*. ^[10]^. *E. coli* strain 25922 (ATCC) was used as the reference strain.

### DNA isolation and polymerase chain reaction assays

Enteropathogenic *E. coli* DNA extraction done according to Cheng and Jiang ^[11]^ method. DNA templates subjected to PCR using six sets (F and R) of primers targeting *bla* genes listed in Table (1).

The reaction mixture moreover contain GoTaq® Green Master Mix, X2 (Promega M 7122; Promega Corporation, USA) and according to Promega procedure, the reaction mixtures were prepared in 0.2 ml eppendorf tube with 25 μl reaction volumes. PCR performed with PCR system (GeneAmp PCR system 9700; Applied Biosystem, Singapore). The PCR amplification conditions performed with a thermal cycler were specific to each single primer set (Table 1) depending on their reference procedure. The amplified PCR products detected by agarose gel electrophoresis and visualized by staining with ethidium bromide by UV-transilluminator.

**Table 1:**
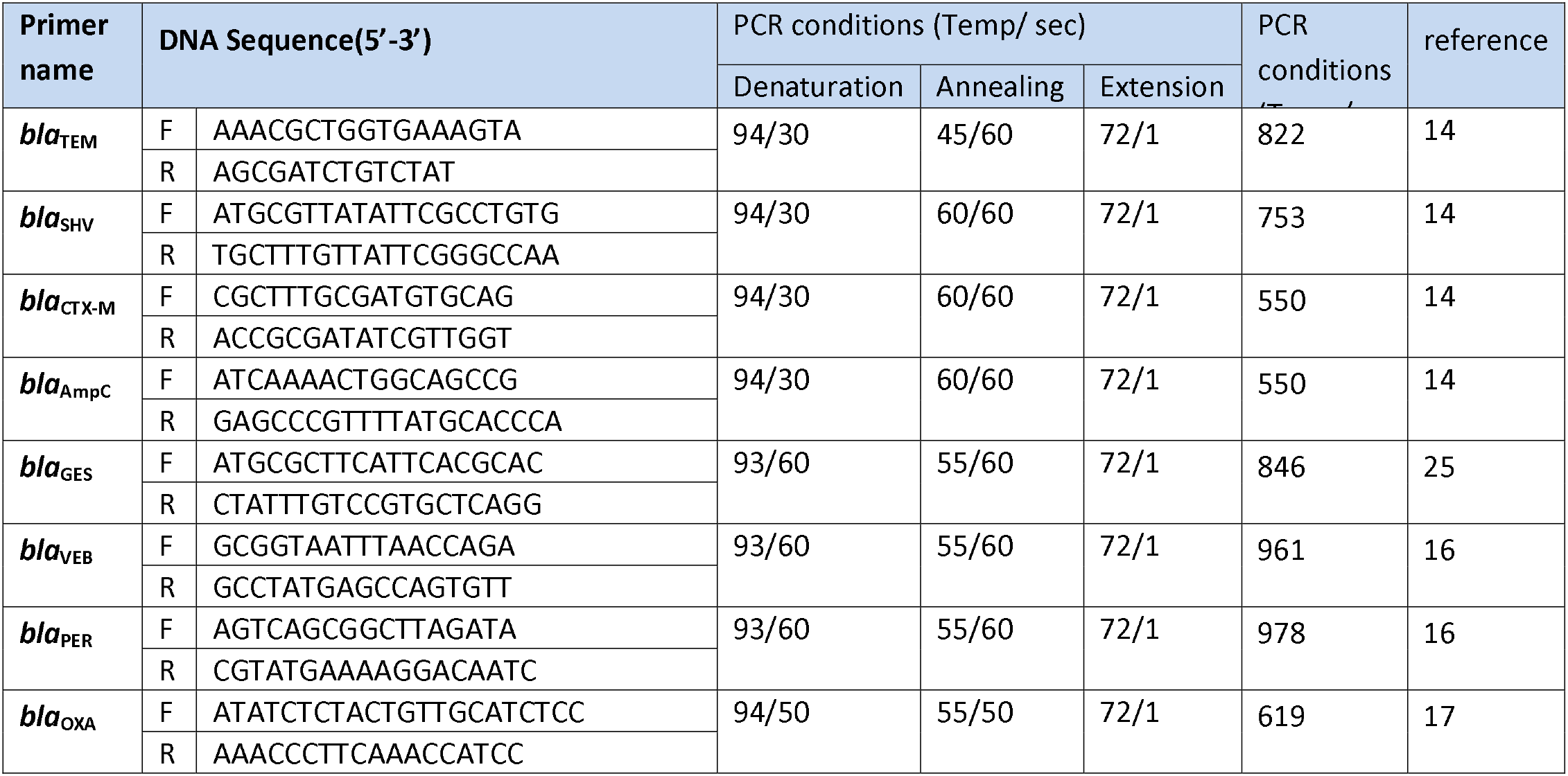
Primer sequence for detection β-lactam resistance genes using PCR assay.

## Results

Of the 656 diarrheal stools analyzed, 535 specimens yielded biochemically confirmed *E. coli* isolates, only 22 (3.4%) agglutinated with EPEC antisera. The *E. coli* serotypes identified shown in Table (2). Twenty (90.9%) isolates were resistant to a minimum of 3 classes of antibiotics, to which they were tested, hence the isolates are considered to be multidrug resistant (MDR) ^[12]^. The resistant effect of isolates to carbenicillin amoxicillin piperacillin ticarcillin and amoxicillin/clavulanic acid are different. The rates of resistance were 90.9% for carbenicillin and 86.4% for others. It should be note that the highly diverse resistance rates against cephalosporins ranging from 59.1% to 77.3%. High resistant rate for cefoxitin (90.9%), most isolates were resistant to aztreonam (59.1%). However, all the EPEC isolates were resistant to at least one of the cephlosporins or azetreonam. The most effective β-lactam antibiotics were imipenem and meropenem (100% susceptible). Low rates of resistance to aminoglycosides amikacin, tobramycin and gentamicin were detected in isolates, (0.0%), (4.5%) and (9.1%), respectively, and all were susceptible to the fluoroquinolones tested Table (3).

**Table 2:** Distribution of *bla*-genes in EPEC isolates.

| Isolate<br>Designation | Serotype | *Bla*-gene |  |  |  |  |  |  |  |
| --- | --- | --- | --- | --- | --- | --- | --- | --- | --- |
|  |  | TEM | SHV | CTX-M | OXA | RER | VEB | GES | AmpC |
| E1 | O111:k58(B4) | + | + | + | - | - | - | - | - |
| E2 | O44:k74(L) | + | + | + | - | - | - | - | - |
| E3 | O125:k70(B1)5) | + | + | + | - | - | - | - | - |
| E4 | O55:k59(B5) | - | + | + | - | - | - | - | - |
| E5 | O125:k70(B1)5) | + | + | + | - | - | - | - | - |
| E6 | O44:k74(L) | + | + | + | - | - | - | - | - |
| E7 | O128:k67(B1)2) | + | + | + | + | - | - | - | - |
| E8 | O127:k63(B8) | - | + | + | - | - | - | - | - |
| E9 | O44:k74(L) | + | + | - | - | - | - | - | - |
| E10 | O111:k58(B4) | + | + | + | - | - | - | - | - |
| E11 | O26:k60 (B6) | + | + | + | - | - | - | - | - |
| E12 | O111:k58(B4) | + | + | - | - | - | - | - | + |
| E13 | O111:k58(B4) | + | + | + | - | - | - | - | - |
| E14 | O26:k60(B6) | - | + | - | - | - | - | - | - |
| E15 | O55:K59(B5) | + | + | + | - | - | - | - | - |
| E16 | O55:K59(B5) | + | + | + | - | - | - | - | - |
| E17 | O55:K59(B5) | + | + | + | - | - | - | - | - |
| E18 | UT (polyvalent 4) | + | + | + | - | - | - | - | - |
| E19 | UT (polyvalent 2) | - | + | - | - | - | - | - | + |
| E20 | O114:k90(B) | + | + | + | - | - | - | - | - |
| E21 | O55:K59(B5) | + | + | - | - | - | - | - | - |
| E22 | O26:K60(B6) | + | + | + | - | - | - | - | - |
| **Sum** |  | **18** | **22** | **17** | **1** | **0** | **0** | **0** | **2** |

**Table 3:** Frequency of antibiotics susceptibility of 22 EPEC isolates in Najaf.

| Antibiotics | EPEC isolates- |  |  |
| --- | --- | --- | --- |
|  | Sensitive | Intermediate | Resistance |
| Amoxycillin | 3(13.6%) | - | 19(86.4%) |
| Piperacillin | 1(4.5%) | 2(9.1%) | 19(86.4%) |
| Carbenicillin | 1(4.5%) | 1(4.5%) | 20(91%) |
| Ticarcillin | 3(13.6%) | - | 19(86.4%) |
| Amoxyclav | 3(13.6%) | - | 19(86.4%) |
| Cefotaxime | 6(68.2) | 1(4.5%) | 15(27.3%) |
| Ceftazidime | 4(18.2%) | 2(9.1%) | 16(72.7%) |
| Ceftriaxone | 9(59.1%) | - | 13(40.9%) |
| Cefepime | 4(18.2%) | 1(4.5%) | 17(77.3%) |
| Cefoxitin | 1(4.5%) | 1(4.5%) | 20(91%) |
| Aztreonam | 9(40.9%) | - | 13(59.1%) |
| Imipenem | 22(100%) | - | - |
| Meropenem | 22(100%) | - | - |
| Amikacin | 22(100%) | - | - |
| Tobramycin | 21(95.5%) | - | 1(4.5%) |
| Gentamicin | 10(45.5%) | 10(45.5%) | 2(9.1%) |
| Ciprofloxacin | 22(100%) | - | - |
| Levofloxacin | 22(100%) | - | - |
| Trimethoprim | 5(22.7%) | 1(4.5%) | 16(72.7%) |
| Tetracyclin | 11(50%) | - | 11(50%) |

All EPEC isolates were screened for ESBL production using DDST; results showed 3 (13.6%) isolates were exhibited zones enhancement with clavulanic acid, confirming their ESBL production. The results confirmed that 20 isolates yield cefoxitin zone diameter less than 18mm (screen resistance). These resistance isolates suspected as AmpC β-lactamase producers. The cefoxitin resistance isolates were further confirmed by the MTDT, a clear distortion of the zone of inhibition of cefoxitin indicating strong AmpC producer was observed in 4 (20.0%) of the cefoxitin resistant isolates.

Table (2) shows the distribution of β-lactamases determined by the consistent results of primers specific PCR. All isolates carried at least one type of *bla* genes. The most commonly identified ESBL gene was *bla*_SHV_ type (100%), while 18 (81.8%) isolates yielded amplification products with TEM-PCR specific primers. The presence of the *bla*_CTX-M_ gene detected in 17 (77.3%) isolates. Only one isolate (4.54%) carried *bla*_OXA_ gene, and 2 (9.1%) isolates were amplified with *bla*_AmpC_ primers.

## Discussion

A variety of antibiotics have been used to treat infection caused by EPEC and have proved useful in many cases, but multiple antibiotic resistances are common among EPEC. Many strains of EPEC known to harbor mobile elements that encode antibiotic resistance and can be transfer among themselves or to other bacterial species to establish multiple antibiotic resistances ^[17]^.

In this study, the MDR rate of EPEC isolates evaluated against common antibiotics. Our results showed that MDR rate in EPEC was 90.9%. These results confirmed data reported by other authors, indicating that EPEC are frequently and increasingly demonstrating multiple resistances to the antibiotics ^[1]^. The high occurrence of MDR isolates of EPEC may be due to the widespread use of antibiotics in Najaf.

Our study revealed that all the EPEC isolates were resistant to at least one of the cephalosporin and azetreonam antibiotics. The high rates of resistance might be as markers for the production of ESBLs by these isolates. However, there are very few reports of ESBL production by diarrheagenic *E. coli* in the Middle East ^[17]^. In Najaf, children with invasive diarrhea might be treat with third-generation cephalosporin, in view of the fact that the majority of EPEC isolates in this study were resistant to third-generation cephalosporins.

The study found that the frequency of ESBL-producing isolates was lower than a study accomplished in Hilla by Almohana et al. ^[19]^ who indicates that ESBL production was confirmed in 36/82 (43.9%) of *E. coli*. There is a number of instances whereby the screening tests for ESBLs are positive but the confirmatory tests negative or indeterminate ^[20]^. Nonetheless, in this investigation not all screened positive EPEC isolates found to be ESBL producers. On the other hand, 86.4% of EPEC found to be resistance to amoxicillin/clavulanic acid, since clavulanic acid inhibits the ESBLs, reducing the level of resistance to the cephalosporins and thereby increasing the zone of inhibition for the disk diffusion tests. Resistance against β-lactamase inhibitors occurs mainly by several mechanisms: hyperproduction of β-lactamases, production of β-lactamases resistant to inhibitors, and chromosomal cephalosporinases ^[21]^. In this study, DDST is a trustworthy, suitable and reasonably priced method of screening for ESBLs. However, this test can lack sensitivity because of problems of optimal disk spacing, the inability of clavulanate to inhibit all ESBL, the inability of test to detect ESBL in isolates that also produce chromosomal cephalosporinase and the loss of clavulanate disk potency during storage^[22]^.

Our results revealed that 81.8% of the isolates yielded amplification products with TEM-PCR specific primers. In Portugal, *bla*_TEM_ gene identified in 85% isolates ^[23]^. We found that all EPEC isolates harbored *bla*_SHV_ gene. Additionally, 77.3% of the isolates harbored a *bla*_CTX-M_ gene. CTX-M constitutes a novel and rapidly growing type of plasmid-mediated ESBLs that is currently replacing mutant TEM or SHV ESBL families and which is much greater penetration into *E. coli*. In the Middle East area, reports from Lebanon and Kuwait pointed out that CTX-M is the predominant ESBL in *E. coli*^[25]^. Even though this may be due to the different antibiotics policies exist in various hospitals with excessive use of third generation cephalosporins. However, some risk factors of the acquisition of *bla*_CTX-M_ gene may be pressure from the surroundings by antibiotics. Results also revealed that only one isolates carried *bla*_OXA_ gene.

Although, cefoxitin is not use in treatment of bacterial infections in Najaf hospitals, present investigation showed that 90.9% of EPEC isolates were cefoxitin resistant. The frequency of cefoxitin resistance in our study was higher than previously recorded in Iraq (44.4%) by Almohana^[19]^. The study revealed that AmpC β-lactamase production confirmed by MTDT in 40% isolates. A limitation of methods used to detect the AmpC enzyme is that an increasing number of clinical isolates that have multiple β-lactamases, which in turn can make inhibition patterns complex and difficult to interpret ^[14]^. There is no standardized method (such as synergy test for ESBL) to easily detect AmpC enzymes. Particularly, it is difficult in *E. coli* to distinguish phenotypically plasmid-mediated AmpC producers from isolates overproducing chromosomal enzymes at high levels. Additionally, a strain with a plasmid-mediated AmpC enzyme can also produce other β-lactamases, such as ESBL, which may complicate the detection of the AmpC phenotype ^[26]^. PCR assay used in this study for the detection of *bla*_AmpC_ gene that proved useful as screening tool to distinguish cefoxitin-resistant non-AmpC producer from cefoxitin-resistant AmpC producer isolate. The PCR assay confirms that only 2 of the 4 AmpC producer isolates previously identified in MTDT carried *bla*_AmpC_ gene.

## Conclusion

It can be concluded that there is a respectively high occurrence of MDR and ESBL-producing EPEC isolates, as well as, high frequency of SHV producing isolates in our country. However, this is the first survey in Najaf hospitals. An important finding for physician in this study as presented in table 1, that the third generation cephalosporins may be not effective for treatment of all cases of EPEC infection in Najaf and the fourth generation cephalosporins and quinolones are the drugs of choices for these cases. This study also states the importance of carrying out sensitivity tests prior to treatment against the use of empirical treatment currently in practice in Najaf.

